# *Campylobacter jejuni* transmission and colonisation in broiler chickens is inhibited by Faecal Microbiota Transplantation

**DOI:** 10.1101/476119

**Authors:** Rachel Gilroy, Gemma Chaloner, Amy Wedley, Lizeth Lacharme-Lora, Sue Jopson, Paul Wigley

**Author notes:** Corresponding author: PW.

## Abstract

**BACKGROUND:** *Campylobacter jejuni,* the most frequent cause of foodborne bacterial infection, is found on around 70% of retail chicken. As such there is a need for effective controls in chicken production. Microbial-based controls such as probiotics are attractive to the poultry industry, but of limited efficacy. Furthermore, as commercially-produced chickens have no maternal contact, their pioneer microbiome is likely to come from the hatchery environment. Early delivery of microbials that lead to a more ‘natural avian’ microbiome may, therefore, improve bird health and reduce susceptibility to *C.jejuni* colonisation.

A faecal microbiota transplant (FMT) was used to transfer a mature cecal microbiome to newly-hatched broiler chicks and its effects on *C.jejuni* challenge assessed. We used both a seeder-bird infection model that mimics natural bird-to-bird infection alongside a direct-challenge model. We used a 16S rRNA gene sequencing-based approach to characterize the transplant material itself alongside changes to the chicken microbiome following FMT.

**RESULTS:** FMT changes the composition of the chicken intestinal microbiome. We observed little change in species richness following FMT compared to untreated samples, but there is an increase in phylogenetic diversity within those species. The most significant difference in the ceca is an increase in *Lactobacilli,* although not a major component of the transplant material, suggesting the FMT results in a change in the intestinal milieu as much as a direct change to the microbiome.

Upon direct challenge, FMT resulted in lower initial intestinal colonisation with *C.jejuni.* More significantly, in a seeder-bird challenge of infection transmission, FMT reduced transmission and intestinal colonisation until common UK retail age of slaughter. In a repeat experiment, transmission was completely blocked following FMT treatment. Delayed FMT administration at 7 days of-age had limited effect on colonisation and transmission.

**CONCLUSIONS:** We show that transfer of a whole mature microbiome to newly-hatched chicks reduces transmission and colonisation of *C.jejuni.* This indicates that modification of the broiler chick microbiome can reduce intestinal colonisation of *C.jejuni* to levels projected to lead to lower the human infection rate. We believe these findings offer a way to identify key taxa or consortia that are effective in reducing *C.jejuni* colonisation and improving broiler gut health.

## BACKGROUND

*Campylobacter jejuni*, a highly motile Gram-negative proteobacteria, is the most frequent cause of human bacterial foodborne gastroenteritis worldwide (1). There are currently estimated to be 9 million cases in the European Union each year, amounting to a vast medical and productivity burden across many of the world’s most developed countries (2). The preparation and consumption of poultry meat continues to be the single largest source of human infection, with over 70% of retail chicken carcasses within the EU showing *C.jejuni* contamination (3). With current intervention strategies aimed at reducing *C.jejuni* burden within the commercial broiler (meat-producing) chicken showing limited success, a pragmatic means of large-scale on-farm control continues to be a key goal (4). The need to develop controls within poultry production without the use of antimicrobials is a public health priority. However, unlike *Salmonella*, where vaccination has proved successful, the nature of both the pathogen and host response to *Campylobacter* in the chicken make the development of vaccines challenging (3). With the gut microbiome acting as the immediate biological barrier against *C.jejuni*, its manipulation could play a key role in its reduction and control in chicken meat production.

Manipulation of the microbiota in livestock has a long history (5). Animal husbandry practices including transfaunication, the transfer or rumen content between cattle and the use of dietary products, particularly probiotics and microbial products in poultry, to manipulate or modify animal intestinal microbiomes to improve health, productivity and wellbeing have long been used (6). Early work using cultured avian intestinal flora to reduce *Salmonella* colonisation in chicks by Rantala & Nurmi (1973) was the forerunner of many subsequent studies on probiotics but the basis of how any manipulation of the microbiome is effective in reducing pathogen load in chickens remains unclear, though broadly there would appear to be two main possibilities. Firstly, any preparation may have a competitive exclusion (CE) effect, originally an ecological term, based around competition for a niche and resources. We also now understand that intestinal tract bacteria such as *Firmicutes* produce metabolites such as butyrates that can inhibit the growth of proteobacteria (7). Secondly, probiotics and microflora preparations may drive immune development and immunity in the gut helping limit pathogen colonisation (8).

Attempts at reproducing and improving such probiotic efficacy in reducing the colonisation of the avian gastrointestinal tract (GIT) with *C.jejuni* has been of largely empirical nature with little evidence of a practical industrial role (8,9). With oral probiotics, doses often provide a relatively low magnitude of microorganisms compared to that found within the native microbiome making repeated administration necessary (5). Although commonly derived from the avian intestinal tract, environmental adaptation during *ex vivo* culture may limit beneficial impact of probiotic formulations which may no longer display the same phenotype as when in the gut (5). Consideration of the gut microbiota as an entire system as opposed to the sum of individual entities offers potential for a more viable solution to *C.jejuni* control. The use of more complex but undefined microbiota preparations such as Aviguard® or Broilact® have been increasingly adopted in Europe, though these are cultured products that are unlikely to contain the full complement of species or genera found in the healthy microbiome. However, their undefined nature precludes their use in many countries such as the United States.

The introduction of a complete, stable gut microbiome from a healthy donor into a recipient through a Faecal Microbiota Transplant (FMT) has recently been incorporated into the therapeutic treatment of an array of known and idiopathic conditions (5). Perhaps the best described and most effective clinical use of FMT in human medicine is to treat recalcitrant *Clostridium difficile* infection (CDI), a result of dysbiosis stemming from antibiotic use, is one of the most notable examples of current therapeutic benefit. A study by Aas et al (2003) presented a FMT treatment success rate exceeding 90% within trial evaluable patients, such findings being reproducible throughout considerable further research (10,11). Although the scientific rationale behind its efficacy remains somewhat elusive, the undoubtable success of FMT in the treatment of CDI warrants further indication of multiple applications beyond current practice. While use of FMT is becoming progressively disseminated throughout human clinical practice, FMT in a modern sense has not yet been adopted into livestock. Here we transfer a faecal, or more strictly a cecal, microbiome transfer from eight-week old animals to newly hatched chicks and using challenge models assess the effect of FMT on host susceptibility to *C.jejuni* infection and its transmission. We determine how FMT alters the microbiome through 16S rRNA gene-based sequencing, with a view towards a rational approach of determining individual bacterial taxa or consortia that offer protection against *C.jejuni* infection.

## RESULTS

### Early faecal transplant has significant impact on *C.jejuni* colonization of the Ceca and ileum following experimental seeder infection

Early faecal microbiota transplantation significantly reduced *C.jejuni* M1 load in both the ceca and ileum following experimental seeder infection of broiler chickens (Table 1). Birds receiving FMT showed a significant reduction in *C.jejuni* load within the ileum when compared to Hatchery control birds in both seeder Experiments 1 and 2 (P=0.0007; P=0.0451). The impact was even greater in reducing colonisation further along the tract in the ceca (P<0.0001; P<0.0001) (Figure 1). Using direct challenge, rather than seeder-bird challenge, colonisation was significantly lower in the ceca and ileum of birds given FMT (P=0.0035; P=0.0152) at 4dpi (days post-infection). However, at 10dpi there was no significant difference between the treatment populations indicating initial inhibition but not prevention of colonisation (Figure S3 in Supplementary Material).

**Table 1.** Levels of *C.jejuni* M1 in ceca and ileum of broiler chickens under experimental conditions according to Experiment 1 and Experiment 2 protocols.

|  | EXPERIMENT 1 |  | EXPERIMENT 2 |  |
| --- | --- | --- | --- | --- |
|  | Median *C.jejuni* M1 content at 14 dpi (CFU.g<sup>-1</sup>) |  | Median *C.jejuni* M1 content at 12 dpi (CFU.g<sup>-1</sup>) |  |
|  | [IQR] |  | [IQR] |  |
|  | Ceca | Ileum | Ceca | Ileum |
| FMT | 4.40 X 10<sup>4</sup> [5.80 X 10<sup>7</sup>] | 0.00 [0.00] | 0.00 [0.00] | 0.00 [0.00] |
| Ext. Hatchery Control | 1.70 X 10<sup>11</sup> [1.10 X 10<sup>11</sup>] | 4.00 X 10<sup>3</sup> [2.42 X 10<sup>4</sup>] | 3.78 X 10<sup>8</sup> [1.56 X 10<sup>9</sup>] | 0.00 [6.15 X 10<sup>3</sup>] |

**Figure 1.**
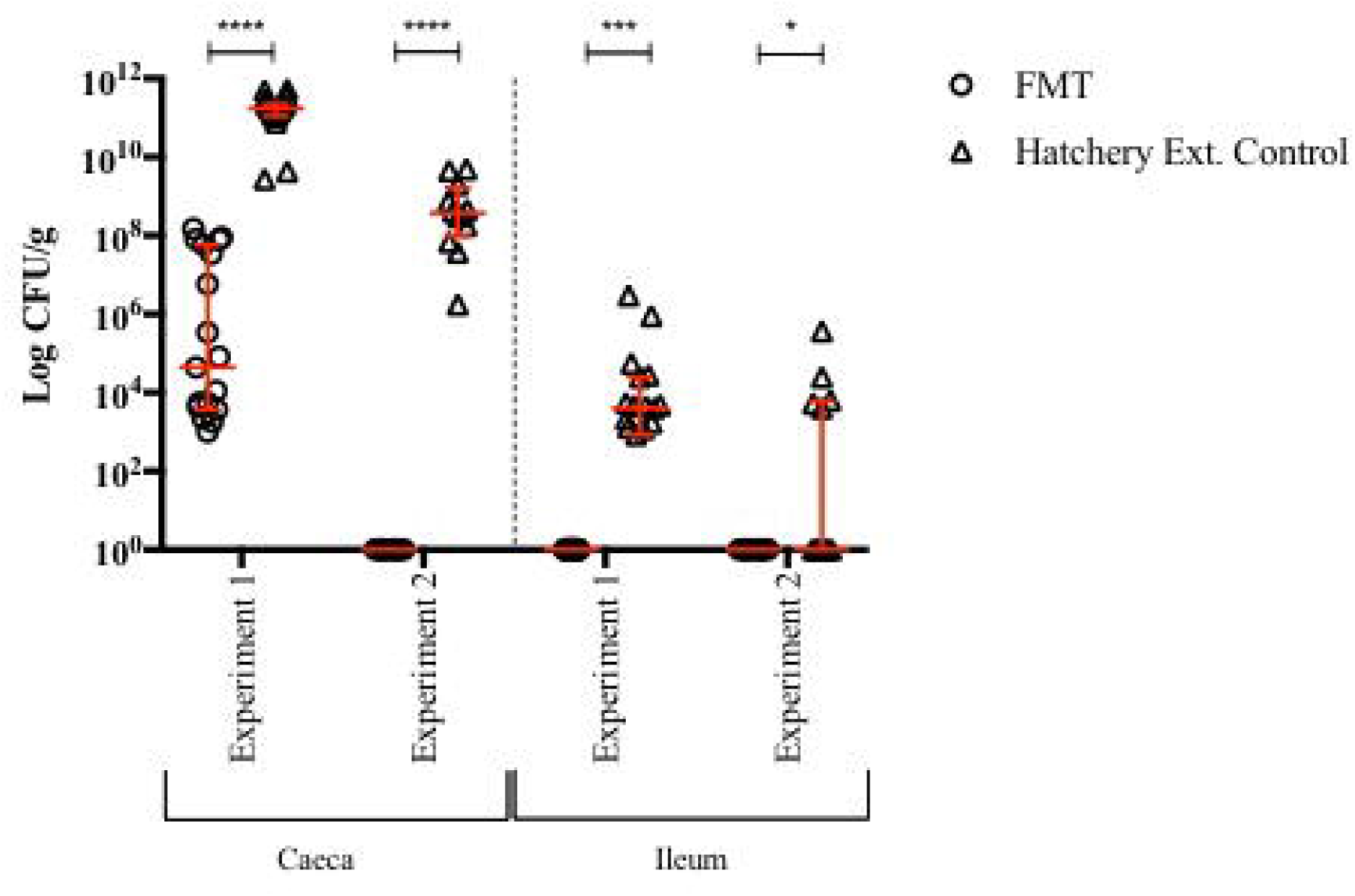
Levels of Cjejuni Ml in the ceca and ileum of broiler chickens grown under cxpcrimcnlal conditions hisedon protocols for Experiment 1 and Experiment 2. Each symbol represents C.jejunicolonisation loud for anintitiiiial sample. Results aie also express'd as median values and associated IQR with significance determined using Man whitney-U analysis. Levels of significance given are at *p<0.06** p<0.01. ***p<0.001. ****p<0.0001.

### Transmission of *C.jejuni* within an experimental broiler flock is delayed by early faecal microbiota transplantation

Between 2dpi and 14dpi cloacal swabs were used to determine the dynamics of *C.jejuni* infection within each population of birds. The kinetics of transmission were considerably slowed within the FMT group compared to that of the untreated hatchery group (Figure 2). Experiment 1 showed 18/19 hatchery birds were shedding *C.jejuni* at 5dpi, whereas there was no detected shedding within the FMT population. All 19/19 birds were shedding *C.jejuni* M1 by 8 dpi within the Hatchery external control population and all birds continued to shed until 12 dpi. There was no bacterial shedding within the FMT population until 12dpi, with 4/19 birds shedding *C.jejuni* by this time-point. The transmission dynamics within experiment 1 were similar within experiment 2, with no *C.jejuni* shedding detected in the FMT group during the course of the trial. Shedding was detected within the Hatchery group from 3dpi and by 10dpi 11/12 birds were shedding *C.jejuni* (Figure 2).

**Figure 2.**
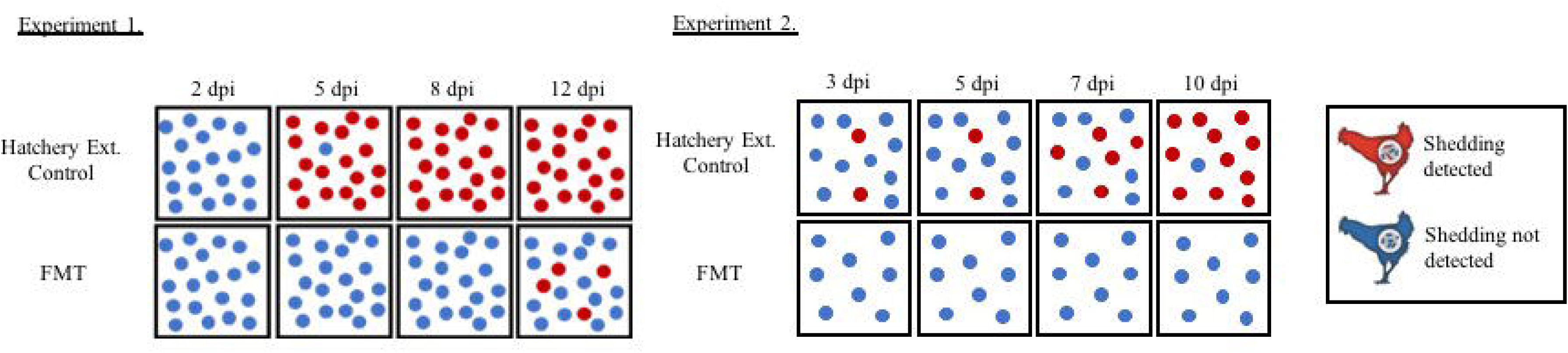
*C.jejuni* M1 transmission within broiler groups of Experiment land Experiment 2 determined through cloacal swabbing. Red shapes show birds where bacterial shedding was detected while blue shapes show groups with no bacterial shedding.

### Faecal microbiota transplantation administration at 7 days of age has no significant impact on *C.jejuni* GIT colonisation and transmission

Swabs taken between 3dpi and 10dpi in experiment 3 showed a slight delay in transmission of *C.jejuni* within the FMT population compared to that of the Hatchery, however this was not sustained. At 3dpi 1/15 birds was shedding *C.jejuni* within the group of birds given the FMT while this number was 3/17 within the external control hatchery group. However, by 5dpi the difference in level of shedding between the two groups was negligible, with this relationship continuing until swabbing at 10dpi whereby all birds in both groups were shedding *C.jejuni*. There was no significant reduction in final levels of *C.jejuni* colonisation within either the ceca or the ileum (P=0.2403; P=0.1268) at 12dpi of experiment 3 populations.

### Extra intestinal spread of *C.jejuni* may be reduced following early faecal transplant administration

At post-mortem examination, extra intestinal *C.jejuni* colonisation was present in all 3 experimental trials. Experiment 1 showed *C.jejuni* within the liver tissues of 2/19 Hatchery birds and 1/19 birds given FMT. This result was similar in experiment 2, with *C.jejuni* present in 2/12 liver and 1/12 spleen samples from Hatchery birds. No *C.jejuni* colonisation was detectable within the FMT population of this experiment. Both Hatchery and FMT populations of experiment 3 showed high liver (>75%) and spleen (>18%) infection. These results further confirm the invasive ability of *C.jejuni* M1 to spread beyond the GIT (Humphrey *et al.*, 2014).

### Cecal microbiota contains similar bacterial species richness following FMT but phylogenetic diversity within those species is increased

Pre-infection microbiota samples taken 7 days post-hatch from Internal control, Hatchery and FMT populations of experiment 1 were sequenced using Illumina MiSeq sequencing protocols. Targets amplification of the hypervariable V3/4 region of the 16S rRNA gene was used and amplified reads clustered into Operational Taxonomic Units (OTUs) based on 97% similarity. The number of unique OTUs observed within each sample following filtering ranged from 538 to 1833 with a total of 2613 unique OTUs observed across all tested samples. A core microbiome of 1874 shared phylotypes was present between different treatment groups. When directly comparing all treatment groups, FMT and Hatchery external control groups showed fewest shared observed OTU’s (Figure 3a). Alpha rarefaction was performed on all samples, with the depth based on the median count of sequences found per sample. Faith’s phylogenetic diversity and Chao1 index were used as measures of community evolutionary distance and predicted species richness respectively within samples (Figure 3b, 3c).

**Figure 3.**
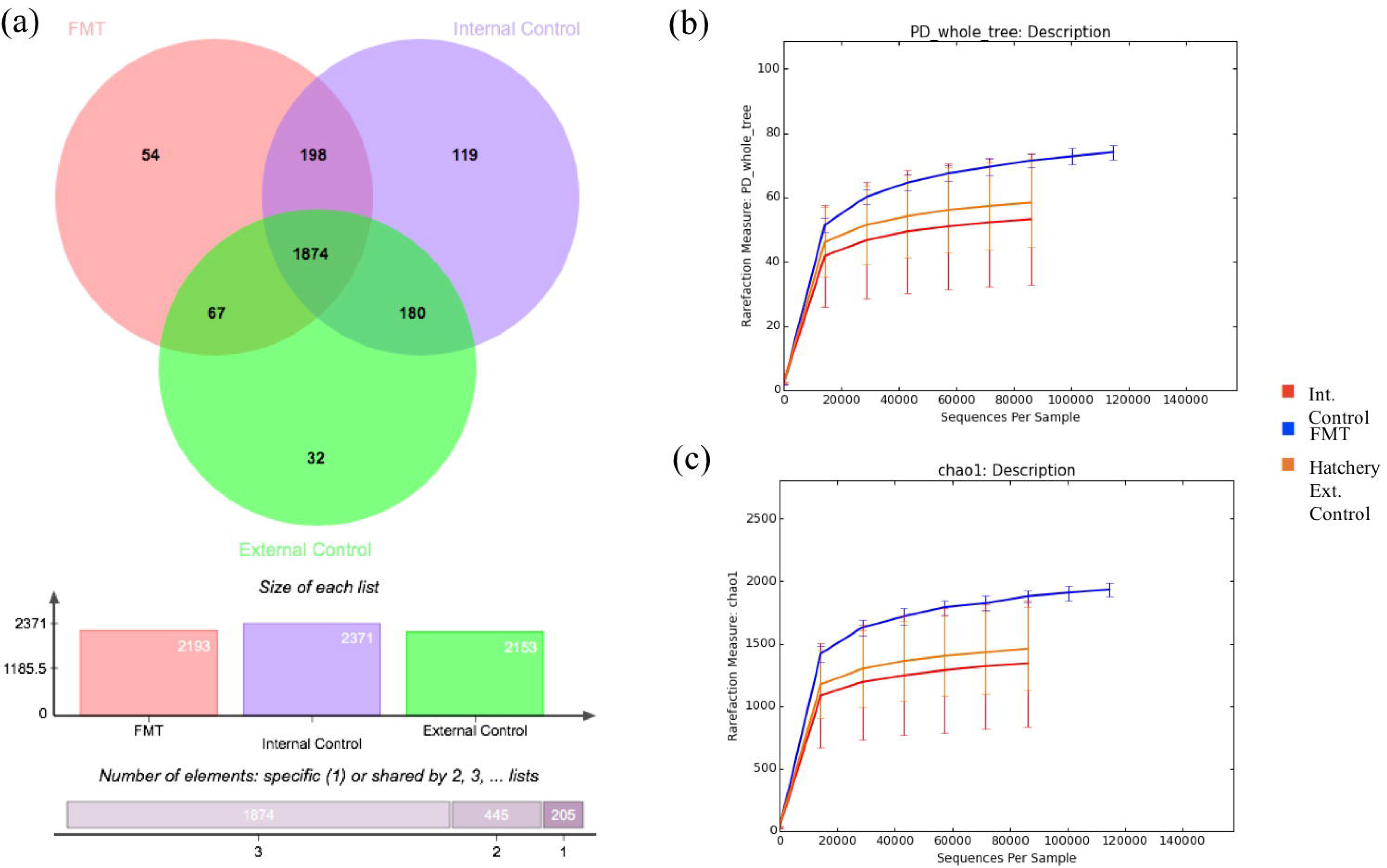
Venn diagram showing the shared gut microbiome (OUT.s) between the 3 treatment groups of Experiment 1 pre *C.jejuni* infection (a). Alpha rarefaction curves were generated based on the OLTPs identified using 97% sequence similarities for FMT, Hatchery External control and Internal control groups using a measure of Faith's phylogenetic diversity (b) and Chao 1 (c).

Shannon diversity index was used to characterize this taxonomic diversity within each sample, with FMT and Internal control populations showing significant difference in diversity (P=0.0357). FMT treated population may therefore show a similar microbiome species richness compared to the Hatchery and Internal control populations, but the phylogenetic diversity seen within the phylotypes present is greater.

### The taxonomic composition of FMT microbiota was distinct to that of Hatchery and Internal control populations

To identify possible variations in the community structure of the gut microbiota by treatment, we calculated the beta-diversity of the samples using PCoA transformation of weighted UniFrac matrixes. The FMT population showed significant clustering and distinct spatial separation from the Hatchery external control and Internal control populations using ADONIS analysis (P=0.001, R^2^=0.361) (Figure 4). These data suggest that the provision of a faecal microbiota transplant immediately post-hatch alters the overall composition of the gut microbial community compared to those not having received treatment.

**Figure 4.**
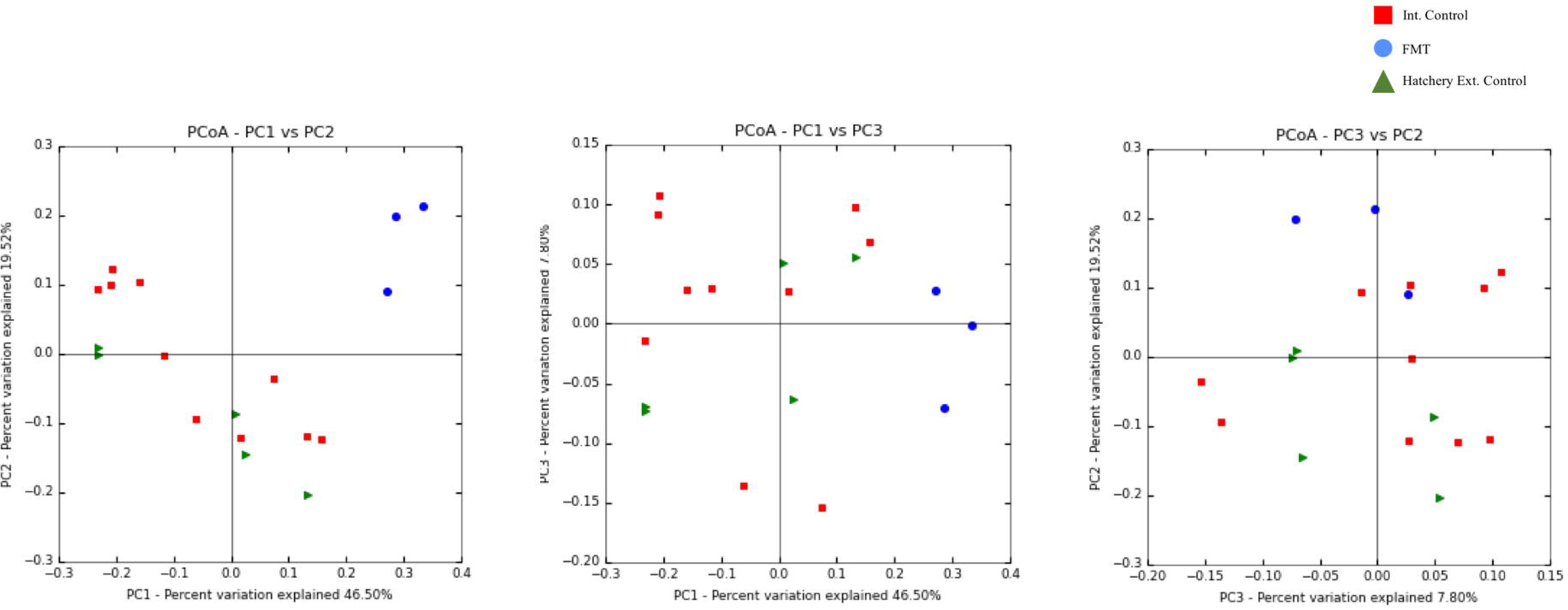
Principal coordinate analysis (PCoA) plot based on weighted UniFrac distances for FMT, Hatchery control and Internal control experimental populations from Experiment 1. Points represent individual samples in the data set. ADONIS testing revealed a significant clustering of samples (P<0.05). PCI and PC2 explain >65% of the total variance within the dataset. Red data points: Internal control; Blue data points: FMT; Green data points: Hatchery external control.

To examine differential representation of taxa between our sample groups, we compared relative abundance at multiple taxonomic levels. The 2613 phylotypes (OTUs) identified were classified into four known and one unknown phyla, with *Firmicutes* predominating all samples at a relative abundance of >90%. Further taxonomic classification at order level showed FMT samples had an average 4.50 fold increase in relative abundance of *Lactobacillales* and an average 1.78 fold decrease in relative abundance of *Clostridiales* compared to both Internal control and Hatchery external control treatment populations (Figure 5, Figure 6).

**Figure 5.**
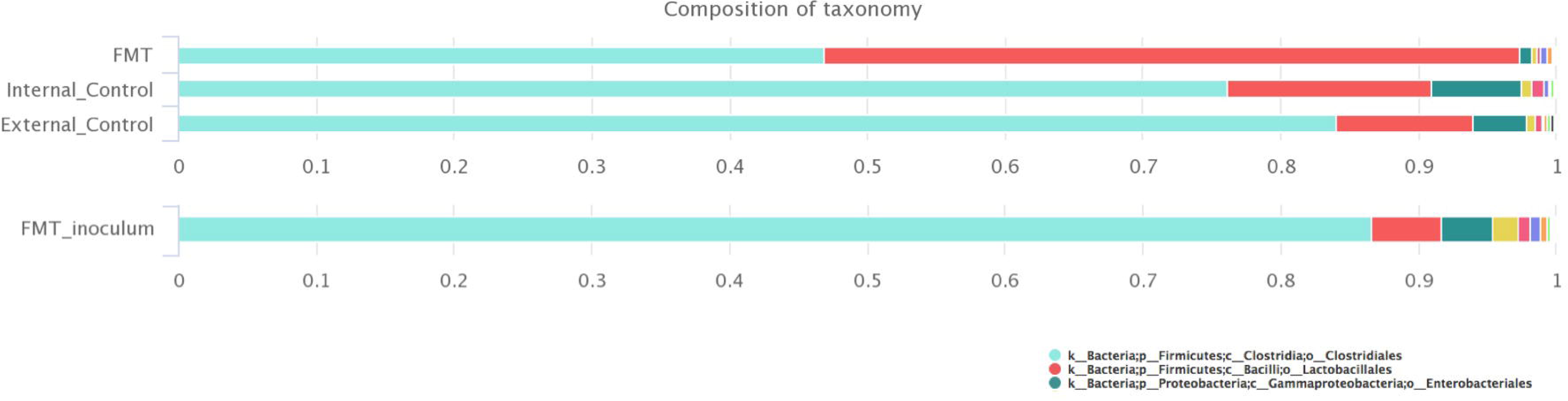
Microbial composition of samples from FMT, Hatchery external control and Internal control treatment populations following 16S rRNA gene sequencing. Included is detail on the composition of FMT inoculum material given to all birds within FMT treatment populations. Composition is displayed as relative percentage abundance of the common taxa (Order) within that treatment group. The 3 most abundant taxonomic groups within treatment samples and inoculum are listed.

**Figure 6.**
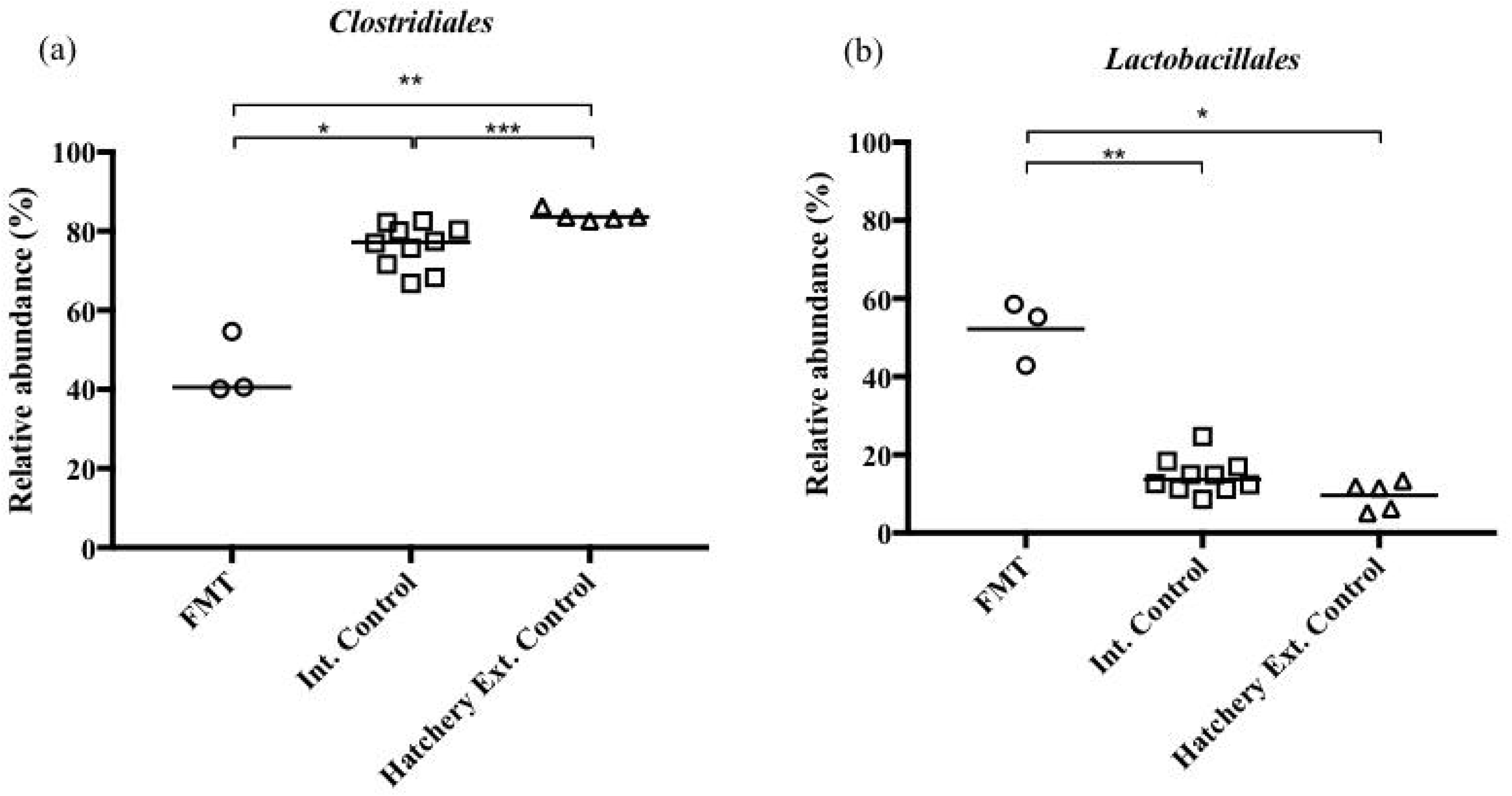
Relative percentage abundance of Clostridialcs (a) and Lactobacillales (b) taxa within samples from each treatment group following 16S rRNA gene sequencing of samples from Experiment 1. Each symbol represents relative abundance within an individual sample. Results are also expressed as median values with significance determined using Mann Whitney-U analysis. Levels of significance are given at *p<0.05, **p<0.01, ***p<0.001, ****p<0.0001.

### *Clostridiales* formed the major taxa within administered FMT inoculum

Samples of FMT inoculum were taken for V3/V4 16S rRNA gene sequencing using Illumina MiSeq sequencing protocols as mentioned previously. Comparing relative abundance of different taxa at multiple taxonomic levels found *Firmicutes* as the predominant phyla at >95% abundance, as with samples taken from our treatment groups. Further classification identified *Clostridiales* as having an abundance >86%, being the core taxa with samples of administered FMT inoculum (Figure 5). The number of observed OTUs within the FMT inoculum itself was the same as that culminated within the treatment group samples.

## DISCUSSION

Here we show that transplanting a whole developed intestinal microbiome from older birds to newly-hatched chicks leads to the long-term modification of the intestinal microbiome, which decreases experimental transmission of *C.jejuni* within a flock of broiler chickens. We propose that early administration of a complex microbiota offers clear potential to reduce *C.jejuni* infection in chicken meat production. Whilst a transfer of a whole microbiota may be impractical in commercial production where billions of birds are reared worldwide each year, it does represent a significant tool in finding consortia of microorganisms that protect against infection in a rational manner.

One fundamental aspect allowing for the potential success of microbiota-based interventions in commercial chicken production is the very nature of large-scale poultry meat production itself. Unlike other livestock species, commercially-produced poultry have no contact with their mothers and do not acquire a pioneer microbiome through maternal transfer, but through the environment of the hatchery (8). This leads to a lack of early diversity and a potentially ‘humanised’ microbiome via hatchery workers (8). Provision of an early, more ‘avian’ microbiome could help drive gut and immunological development leading to a healthier gut and an animal less able to be colonized by *Campylobacter*.

While no statistically significant differences in species richness were found between treatment groups, it is notable that in terms of the cecal microbiota taxa observed there were marked differences between treatment groups regarding phylogenetic diversity. It was found that chicks hatched and reared within our poultry unit as internal controls showed closer phylogenetic similarity to hatchery obtained chicks when compared to FMT treated chicks also reared within our unit. As such, it is likely to be the FMT that is contributing most to the stable shift in microbiome composition within the FMT treatment population. Administration of transplant material at a week-of-age had limited impact on transmission and colonization, suggesting that provision of an early and diverse microbiome is important.

It is also interesting that main difference in taxa is the increase in *Lactobacillus* in the transplanted birds, although this taxon does not form a large part of the transplant material (~5%). This suggests that transplant may as much change the intestinal milieu to a more beneficial one, rather than just simply form the basis of the microbiome. The change towards a cecal microbiome rich in *Lactobacilli* is perhaps indicative of this as usually these form a small part of the microbiome in the lower intestinal tract, though are considered beneficial to chicken gut health, being the basis of many probiotics. Moreover, several recent studies have indicated that low levels of *Lactobacilli* in the chicken intestinal tract are associated with an increased load of *Campylobacter*, with links being made to modulation of cytokine gene expression altering immune response or the production of organic acids and anti-campylobacter proteins (12–14). Here, we show higher levels of *Lactobacillus* following transplantation correlate with reduced levels of *C.jejuni* colonization or even exclusion from the ceca following FMT.

As yet, we have not defined the mechanism or mechanisms that reduce *C.jejuni* transmission following cecal transplantation. As discussed previously these are likely to be either competitive exclusion effects or enhanced immune protection. Future work will look assess avian immune response to FMT administration, further characterize microbiome shifts and alternative methods of FMT inoculation that could be utilised on a larger, industrial scale.

## CONCLUDING REMARKS

Together, our data indicate that at-hatch transplantation of an adult microbiome significantly delays *C.jejuni* colonization and transmission at a flock level. The provision of a complete, rather than culturable ‘chicken’ microbiome acts to improve chicken gut health and impede *C.jejuni* in a naturalistic model of infection. We suggest that it is essential that microbiota administration occurs immediately post-hatch to replace the naïve and dynamic chick microbiota with that of a stable ‘chicken’ microbiome. We believe this concept could offer an effective, low-cost control strategy to *C.jejuni* within the poultry industry.

## METHODS

### Bacterial strains and culture conditions

*C.jejuni* M1 was cultured from frozen stocks maintained at −80°C on Colombia blood agar supplemented with 5% defibrinated horse blood (Oxoid, Basingstoke, Hampshire, United Kingdom) for 48h in microaerobic conditions (80% N_2_, 12% CO_2_, 5% O_2_, and 3% H_2_) at 41.5°C. Liquid cultures were grown for 24h in 10ml of Mueller-Hinton broth (MHB) in microaerobic conditions at 41.5°C and adjusted by dilution in fresh MHB to a final concentration required. All microbiological media were purchased from Lab M Ltd. (Heywood, Lancashire, United Kingdom).

### Faecal microbiota preparation

The microbiota was obtained from three *Campylobacter* free, 8 week old Ross 308 birds, reared under bio-secure condition The birds were euthanised before caecal contents were aseptically removed. Caecal contents were then diluted 1:20 in Phosphate Buffered Saline (PBS) solution, filtered through a 25µM filter and stored at −80°C until use.

### Experimental animals

All work was conducted in accordance with United Kingdom (UK) legislation governing experimental animals under project license PPL 40/3652 and was approved by the University of Liverpool ethical review process prior to the award of the licenses. All animals were checked a minimum of twice-daily to ensure their health and welfare. For experiment 1 and 2, embryonated Ross 308 hens’ eggs were obtained from a commercial hatchery and incubated in an automatic roll incubator under standard conditions for hen eggs. Chicks were removed from the incubator post-hatch and an inoculum of FMT was administered to each chick within 4 hours of hatching. A small group (n=10) of hatched chicks were not given any FMT inoculum to act as a separate Internal control group in experiment 1 and used solely for 16S rRNA gene analysis. Age matched, 1 day-old mixed sex chicks of Ross 308 broiler chickens were obtained from the same commercial hatchery and not given the FMT to act as a Hatchery external control group.

Chicks were housed in the University of Liverpool high-biosecurity poultry unit as with (1). At 7 days post-hatch during experiment 1, a small number of chicks from the FMT (n=3) and Hatchery control (n=5), alongside all Internal control (n=10) chicks were culled and microbiota snap frozen for use in 16S rRNA gene sequencing protocols. For further clarification on each experimental protocol see Figure S1&2 in Supplementary Materials.

Prior to experimental infection, all birds were confirmed as *Campylobacter* free by taking cloacal swabs, which were streaked onto selective blood-free agar (modified charcoal-cefoperazone-deoxycholate agar [mCCDA]) supplemented with *Campylobacter* enrichment supplement (SV59; Mast Group, Bootle, Merseyside, United Kingdom) and grown for 48h in microaerobic conditions at 41.5°C.

### EXPERIMENTS 1 & 2. Effect of faecal transplantation on seeder *C.jejuni* infection

At 21 days post-hatch, two birds from both the FMT (Experiment 1 n=19; Experiment 2 n=8) and the Hatchery control (Experiment 1 n=19; Experiment 2 n=12) groups were orally infected with 10^6^ cells of *C.jejuni* M1 in 0.2ml of MHB. Challenge at 21 days of age has previously shown to be a robust model that mimics the situation in the field in the UK, where birds typically become infected at around three weeks of age due to a ‘lag phase’ considered to be a consequence of protection by maternal immunity (4).

At 2, 5, 8 and 12 (Experiment 1) or 3, 5, 7 and 10 (Experiment 2) days post-infection, cloacal swabs of all birds were taken to assess within-group transmission. At 14 dpi (Experiment 1) or 12 dpi (Experiment 2), all birds were culled via cervical dislocation. At post-mortem examination, samples of tissue and gut contents were collected and processed for host bacterial enumeration. The presence/absence of hock marks and/or pododermatitis also was recorded for every bird at post-mortem examination.

### EXPERIMENT 3. Effect of delayed administration of faecal transplant on seeder C.jejuni infection

Age matched, 1 day-old mixed sex chicks of Ross 308 broiler chickens were obtained from a commercial Hatchery and housed as with experiment 1 and 2 in the University of Liverpool high-biosecurity poultry unit. At 7 days post-hatch, a group of birds (n=15) were inoculated with FMT. The remaining birds (n=17) were not given the FMT to act as a control group. At 21 days post-hatch, two birds from both the FMT (n=8) and the Hatchery external control (n=12) groups were orally infected with 10^6^ cells of *C.jejuni* M1 in 0.2ml of MHB. At 3, 5, 7 and 10 days post-infection cloacal swabs of all birds were taken to assess within-group transmission. At 12 days post-infection birds were killed via cervical dislocation. At post-mortem examination, samples of tissue and gut contents were collected and processed for host bacterial enumeration.

### Assessment of *C.jejuni* load

To determine the level of *C.jejuni* intestinal colonisation within each group, cecal and ileal content was collected from individual birds at necroscopy. This was diluted in 9 volumes of maximal recovery diluent (Lab M, Heywood, Lancashire, United Kingdom [MRD]) with further serial 10-fold dilutions being made of each sample in MRD. Using the method as described Miles & Misra (1938), triplicate 20µl spots were plated onto mCCDA agar supplemented with SV59. The plates were incubated under microaerobic conditions at 41.5°C for 48 h, and *Campylobacter* colonies were enumerated to give colony forming units per gram (CFU/g) of cecal and ileal content. Liver and spleen tissue was also biopsied at post-mortem to assess any extra intestinal spread of *C.jejuni* infection. Differences in final colonisation levels between treatment groups were analysed for significance (P<0.05) using Mann-Whitney *U* tests in GraphPad Prism version 7.00 software.

### Assessment of C.jejuni shedding

Cloacal swabbing provided a non-sacrificial method of following *C.jejuni* shedding within individual experimental groups. Cloacal swabs were briefly plated onto mCCDA agar supplemented with SV59. Swabs were then eluted in 2ml modified 5% Exeter broth consisting of 1,100ml nutrient broth, 11ml lysed defibrinated horse blood (Oxoid, Basingstoke, Hampshire, United Kingdom), *Campylobacter* enrichment supplement SV59 (Mast Diagnostics), and *Campylobacter* growth supplement SV61 (Mast Diagnostics). Enriched swabs were then incubated at 41.5 °C for 48h and re-plated onto mCCDA agar and incubated for 48h at 41.5 °C. Plates were assessed for *C.jejuni* positivity.

### DNA Extraction

Cecal and ileal microbiota contents were collected from a random sample of birds in experiment 1 and FMT alongside inoculum samples, snap frozen and stored at −80°C before DNA extraction. Microbial community DNA was extracted from faecal samples using the Qiagen QIAamp^®^ Fast DNA Stool Mini (Qiagen, Hilden, Germany) following the protocol for the Isolation of DNA from Stool for Pathogen Detection. DNA was eluted in 200µl of DNase/RNase Free Water and stored at −20°C until further analysis. Isolated DNA quality and integrity was assessed through 2.0% agarose gel electrophoresis and concentration measured using a Qubit 2.0 Fluorometer (Life Technologies).

### Illumina MiSeq platform sequencing

Extracted DNA was sent for llumina MiSeq sequencing of the V3/V4 hypervariable 16S rRNA gene at the Centre for Genomic Research (University of Liverpool). Sample library preparation and amplification were performed according to the method previously described by D’Amore et al (2016). Prior to data processing, all raw Fastq files were trimmed using Cutadapt version 1.2.1 (15) to remove any Illumina adapter sequences. All reads were subsequently trimmed using Sickle version 1.200 (16) with a minimum window quality score of 20 and any reads containing fewer than 10 base pairs were removed.

### Sequencing data processing

Data processing was performed using the QIIME 1.9.1 pipeline (17). Forward and reverse Fastq reads were joined, filtered and demultiplexed. FASTA file sequences were clustered using USEARCH to create OTUs using a 97% identity threshold (18). Chimeric sequences were removed using UCHIME and the most abundant sequence from each OTU used as representative (19). Taxonomic level for each of these representative sequences was assigned against the Greengenes 16S rRNA gene database (20) and aligned using PyNAST at a minimum identity of 75% (21). OTUs with a number of sequences less than 0.005% of the total number of sequences were removed from further processing (22). A phylogenetic tree was generated using FastTree (23). FMT inoculum samples were then filtered from treatment group samples and analysed separately. Corresponding sequence files and metadata for all samples analysed using 16S rRNA MiSeq sequencing have been deposited in Figshare and are included here as additional files 1 and 2 respectively.

Alpha and Beta diversity metrics were used to assess microbial composition within and between sample groups. Shannon’s Diversity Index, Faith’s Phylogenetic Diversity and Chao1 metrics were all used to assess community richness with Faiths Phylogenetic Diversity further incorporating phylogenetic relationships between features. Alpha rarefaction curves were constructed based on Chao1 and Faith’s Phylogenetic Diversity metrics as a comparison between different treatment groups. A Venn diagram was created using MetaCoMET’s jvenn programme (24) to compare microbiome data from each treatment group with the core microbiome represented within the shared overlapping regions between the circles. A breakdown of the taxa at an Order level contributing to samples within each treatment group were created using MetaCoMET relating to their relative abundance within those samples. Beta diversity principal coordinate analysis (PCoA) estimates were created based on Weighted UniFrac distances (25) to identify similarities between samples under different treatment groups.

## Supporting information

## LIST OF ABBREVIATIONS

EU: European Union
CE: Competitive Exclusion
GIT: Gastrointestinal tract
FMT: Faecal Microbiota Transplant
CDI: Clostridium difficile infection
MHB: Mueller-Hinton broth
PBS: Phosphate Buffered Saline
UK: United Kingdom
dpi: days post-infection
mCCDA: modified charcoal-cefoperazone-deoxycholate agar
MRD: Maximal recovery diluent
CFU/g: colony forming unites per gram
OTU: Operational Taxonomic Unit
PCoA: Principal Coordinate Analysis

## DECLARATIONS

### Ethics approval and consent to participate

All work was performed in accordance with relevant UK legislation of Animal Use (Animals [Scientific Procedures] Act 1986 under project licence PPL 40/3652 which required Ethical Review by the University of Liverpool Animal Welfare and Ethical Review Body prior to its award.

### Consent for publication

Not applicable.

### Availability of data and material

The 16S rRNA sequencing datasets generated and analysed during the current study are available in Figshare as Additional electronic supplementary files.

https://figshare.com/s/469b48d2e022440ce2dc – Additional file 1: MiSeq 16S rRNA sequence files (2.29 GB).

https://figshare.com/s/ae6b0048285f9f2af5fb - Additional file 2: Metadata associated with sample sequences analysed using 16S rRNA sequencing (1.3kB).

### Authors’ contributions

RG and GC led the experimental work and analysis. LL, AW, SJ and PW assisted in the experimental work. RG and PW wrote the manuscript with the assistance of the other authors. PW conceived the study and it was designed in conjunction with RG and GC. All authors approved the final manuscript

## Acknowledgements

We wish to thank Ruth Ryvar, Trevor Jones and Karen Ryan for technical support. We thank Dr Eliza Wolfson (www.lizawolfson.co.uk) for the producing the chicken illustrations.

