## Supplementary figures and images for "*Campylobacter jejuni* transmission and colonisation in broiler chickens is inhibited by Faecal Microbiota Transplantation"

### Supplementary file 1

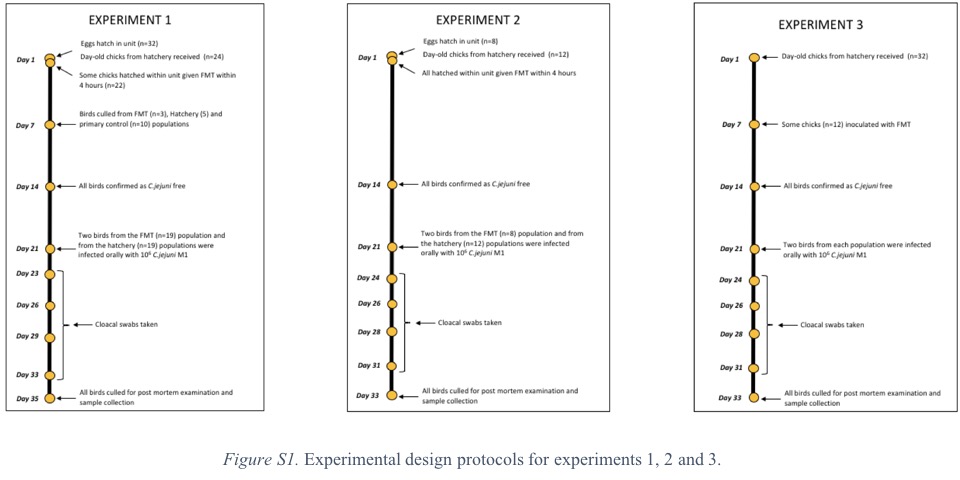

### Supplementary file 2

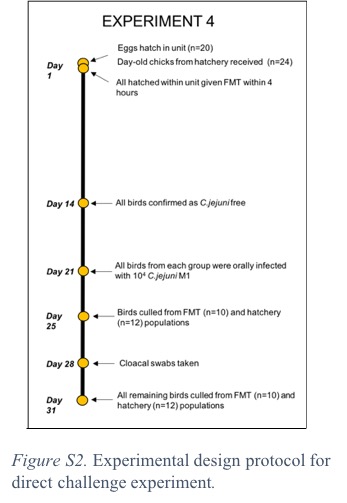

### Supplementary file 3

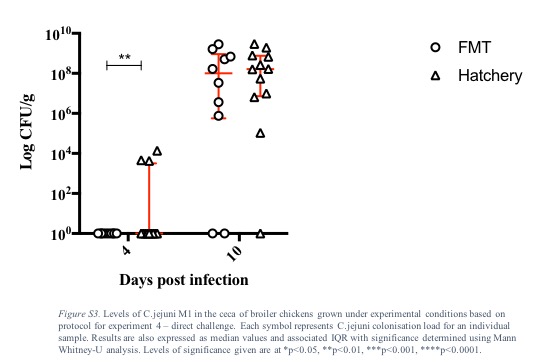
